# Spatiotemporal properties of glutamate input support direction selectivity in the dendrites of retinal starburst amacrine cells

**DOI:** 10.1101/2022.07.12.499686

**Authors:** Prerna Srivastava, Geoff deRosenroll, Benjamin Murphy-Baum, Tracy Michaels, Akihiro Matsumoto, Keisuke Yonehara, Gautam B. Awatramani

## Abstract

The asymmetric summation of kinetically distinct glutamate inputs across the dendrites of retinal “starburst” amacrine cells is proposed to underlie their direction selective properties, but experimentally verifying input kinetics has been a challenge. Here, we used two-photon glutamate sensor (iGluSnFR) imaging to directly measure the input kinetics across individual starburst dendrites. We found that signals measured from proximal dendrites were relatively sustained compared to those measured from distal dendrites. These differences were observed across a range of stimulus sizes and appeared to be shaped mainly by excitatory rather than inhibitory network interactions. Temporal deconvolution analysis suggests that the steady-state vesicle release rate was ∼ 3 times larger at proximal sites compared to distal sites. Using a connectomics-inspired computational model, we demonstrate that input kinetics play an important role in shaping direction selectivity at low stimulus velocities. Together, these results provide direct support for the ‘space-time wiring’ model for direction selectivity.

## Introduction

The radiating dendrites of retinal GABAergic/cholinergic “starburst” amacrine cells (starbursts) are the first points in the visual system to exhibit direction selectivity (Euler et al., 2002). Object motion away from the soma generates large calcium responses in distal starburst dendrites, from where they release GABA and acetylcholine (ACh). By contrast, motion towards the soma evokes weak or no responses at all. Starbursts play a critical role in shaping direction-selective (DS) responses of downstream output ganglion cells (DSGCs) and thus understanding how they compute direction is of principal interest. Decades of intense investigations have identified several mechanisms that underlie direction selectivity in starburst dendrites, although no single mechanism alone may be critically required (reviewed by Murphy-Baum et al., 2021). A model that has garnered recent attention relies on the kinetic properties of distinct sources of glutamatergic input, which is referred to as the ‘space-time wiring’ model for direction selectivity (Greene et al., 2016; Kim et al., 2014). In this study, we sought to evaluate the kinetics of glutamatergic input to the ON starburst dendrites to understand their role in generating direction selectivity.

The space-time wiring model for direction selectivity is inspired by results from connectomic analysis showing that the proximal and distal dendritic regions of ON and OFF type starburst amacrine cells receive synaptic inputs from anatomically distinct types of glutamatergic bipolar cells (BCs). As the axon terminals of these BCs stratify at distinct depths within the inner plexiform layer (IPL; Ding et al., 2016; Greene et al., 2016; Kim et al., 2014)—and in general BC axonal stratification patterns are linked to their kinetic properties (Awatramani and Slaughter, 2000; Baden et al., 2016; Franke et al., 2017; Gaynes et al., 2021; Strauss et al., 2021)—it has been hypothesized that different types of BCs contacting proximal and distal starburst dendrites have distinct response kinetics (Greene et al., 2016; Kim et al., 2014). Specifically, it is predicted that proximal inputs near the starburst soma are mediated by BC types that support tonic patterns of glutamate release, while distal inputs are mediated by BC types that release their vesicles more transiently. This arrangement would result in an optimal input summation along starburst dendrites during centrifugal (soma-to-dendrite) motion, as experimentally noted. The space-time wiring model for direction selectivity is algorithmically similar to classic correlation-type motion detectors described elsewhere in the visual system in both rodents and primates (Lien and Scanziani, 2018; Valois and Cottaris, 1998), as well as the fly optic lobe (Haag et al., 2016; Leong et al., 2016), and thus appears to reflect a core computational principle.

While the space-time wiring model is an attractive model for direction selectivity, there is scant evidence that the kinetics of glutamatergic input varies along starburst dendrites. Direct electrophysiological measurements revealed that BC5s (including types 5i, o and t) and BC7, which make ‘ribbon’ synapses predominantly on the distal and proximal dendrites of ON starbursts, respectively, have similar temporal properties (Ichinose et al., 2014). Regional differences in input kinetics were noted when BC output was measured postsynaptically using voltage-clamp techniques (Fransen and Borghuis, 2017; but see Stincic et al., 2016). However, results from these studies need to be interpreted with caution as the extent to which adequate voltage control is maintained across the length of the thin starburst dendrites during these somatic recordings is unclear, and any voltage escape could severely distort the input kinetics. To this end, genetically-encoded fluorescent glutamate sensors (iGluSnFRs) have provided an alternate way to measure BC output kinetics (Borghuis et al., 2013; Marvin et al., 2013). However, recent imaging studies have found that most BCs of the same polarity (including the BC5s and BC7 types) have similar temporal properties, at least to spatially restricted stimuli that are relevant to local starburst dendritic computations (Franke et al., 2017; Strauss et al., 2021). Finally, recent studies have demonstrated that direction selectivity remains intact under conditions in which all BC input to starbursts is pharmacologically blocked and the starburst network is directly stimulated optogenetically (Hanson et al., 2019; Sethuramanujam et al., 2016). Together, previous results provide little support for the space-time wiring model for direction selectivity.

In the present study, we identify specific stimulus conditions under which stark kinetic differences can be observed in proximal and distal bipolar cell inputs. We did so by directly monitoring input kinetics across starburst dendrites using iGluSnFRs (which we selectively expressed in these cells) across a range of stimulus and pharmacological conditions. In a connectomics-inspired computational model, we used temporal deconvolution to estimate vesicle release dynamics and tested how the specific spatial distributions of kinetically distinct glutamatergic inputs impact starburst direction selectivity. Together, our results indicate that diverse BC kinetics may play a role in shaping direction selectivity, especially in the context of relatively large objects moving slowly across starburst receptive fields.

## Results

### Temporal diversity of glutamate responses along single starburst dendrites

We injected AAVs containing flex-iGluSnFR intravitreally into ChAT-Cre expressing mice. Several days after injection, we sometimes observed a strong, but sparse expression of iGluSnFR in ON-type starburst cells (**Fig. 1 A**). This provided a unique opportunity to visualize glutamate response kinetics across the length of individual dendrites (**Fig. 1 B**). Signals from dendritic regions proximal to the starburst soma were captured in a different optical plane than the more distal dendrites, which are ∼ 5 μm apart (Ding et al., 2016; Greene et al., 2016). Spots of light (200 μm diameter) centered on the imaging field evoked robust iGluSnFR signals throughout the first ∼ 60-80 μm section of starburst dendrites, where glutamatergic BCs are known to make synapses (Ding et al., 2016; Greene et al., 2016). The peak amplitudes of the iGluSnFR signals measured over small regions of interest (ROI; 5 μm X 5 μm) were relatively stable across the length of single starburst dendrites (**Fig. 1C**). However, the sustained phase of the response significantly decreased towards the distal region (**Fig. 1C;** n = 38 ROIs, 2 retinas, 7 dendrites; ΔF/F 0.61 ± 0.24 for proximal, 0.26 ± 0.16 for distal ROIs; *p<0.001, t-test). As a result, the sustained/transient index (STi) computed from the plateau/peak ratio, systematically decreased with increasing distance from the soma (**Fig. 1D**; n = 38 ROIs, 2 retinas, 7 dendrites; STi = 0.35 ± 0.06 for proximal, 0.18 ± 0.06 for distal; *p<0.001, t-test) (Note, STi = 0 indicates a purely transient response with no plateau phase, and STi = 1 indicates a purely sustained response where peak and plateau phases are equal). Together, these results provide the first piece of direct evidence that the kinetics of glutamatergic input varies along starburst dendrites, supporting the ‘space-time’ wiring model for direction selectivity (Kim et al., 2014).

**Fig. 1:**
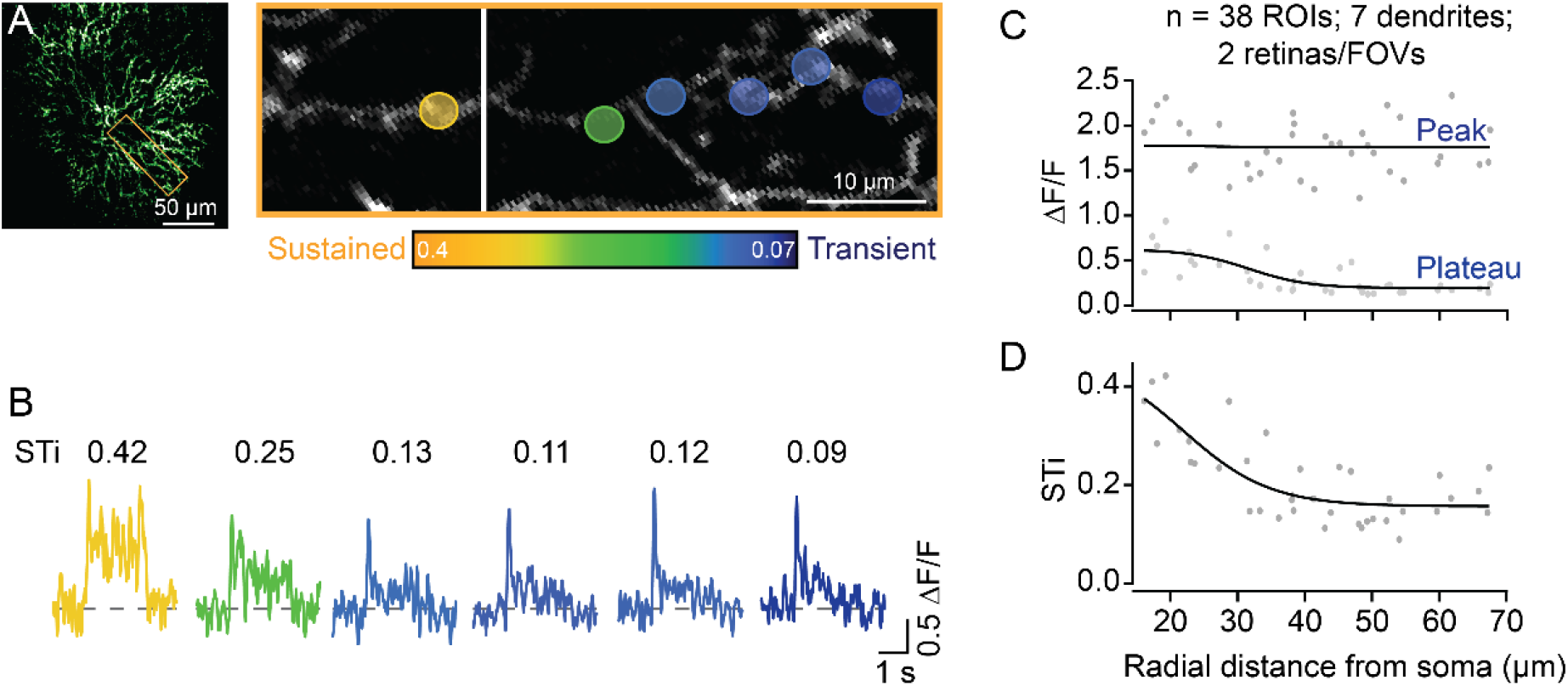
Temporal diversity of inputs across single starburst dendrites revealed by sparse iGluSnFR imaging. **(A)** Two-photon z-stack image (left) of a single ON starburst amacrine cell expressing iGluSnFR. Changes in iGluSnFR fluorescence evoked by a 200 μm spot were measured across the single dendrite (yellow box; left), at two focal planes to capture responses in the proximal and distal regions, and the resulting images were stitched together (right; the vertical white line separates the two fields). (B) Examples of the time-varying iGluSnFR signals (ΔF/F) measured in small dendritic regions of interest (5×5 μm ROIs; shown in A). The responses and ROIs are color-coded according to their STis (color scale bar shown in A). The sustained/transient indices (STis) are indicated above each trace. **(C)** The amplitudes of the peak and plateau responses are plotted as a function of radial distance from the soma. **(D)** STis as a function of radial distance from the soma (n = 38 ROIs from 7 dendrites/2 retinas).

In most experiments, iGluSnFR expression was more widespread, however. In these experiments, individual dendrites leaving the starburst soma were easily visible, but as they dove deeper into the IPL they formed an intricate ‘honeycomb’ mesh. By taking care to lay the retina down flat in the recording chamber, we were able to measure responses from proximal and distal starburst dendrites at separate imaging depths (**Fig. 2A)**. We found that STis were significantly lower in imaging planes that captured distal dendrites (STi = 0.21 ± 0.07; μ ± s.d.) compared to those that captured proximal dendritic responses (STi = 0.34 ± 0.07; μ ± s.d.) (n=247 proximal ROIs; n=563 distal ROIs; 8 retinas; 10 FOVs; **\***p< 0.001, t-test; **Fig. 2A-C**), verifying our initial findings on a larger population level. However, the response latencies and rise times were similar for proximal and distal inputs **(Fig. 2D, E**), indicating that the small differences in axonal path lengths between proximal and distal BCs do not result in significant transmission delays, as previously envisioned (Kim et al., 2014).

**Fig. 2:**
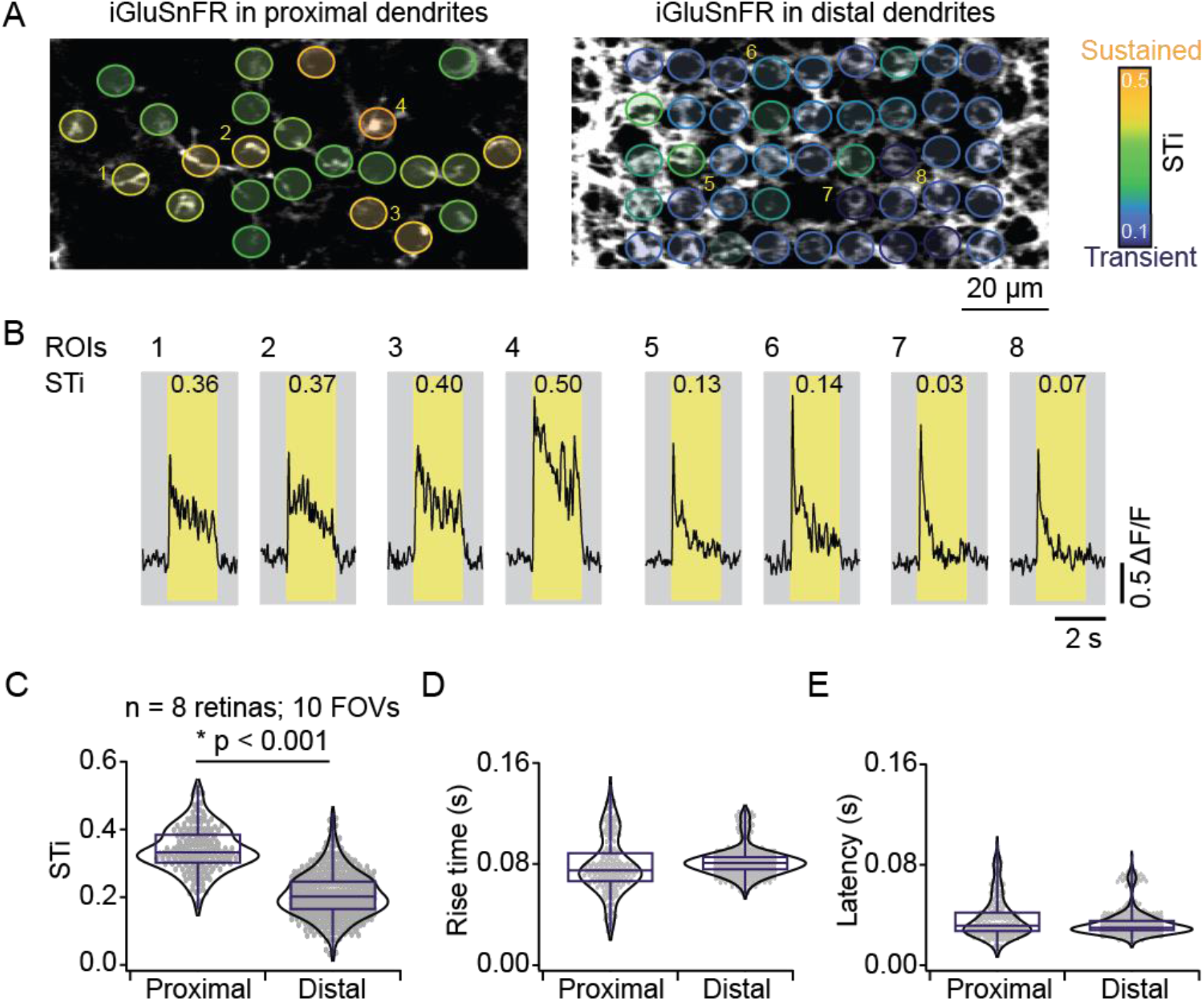
Measuring inputs kinetics in the starburst population. (A) In the left scan field, proximal dendrites arising from the starburst soma expressing iGluSnFR can be visualized in relative isolation. Images taken ∼ 5 μm deeper in the retina reveal the dense honeycomb structure formed by distal starburst dendrites (B) Example iGluSnFR responses evoked by 200 μm static spot extracted for a few ROIs numbered in (A) with their STis indicated on the top. Yellow bands indicate stimulus duration. (C-E) Distribution of STis (C), 80-20 % rise times (D) and latencies (E) in the proximal and distal field of views (FOVs) (n = 10 FOVs, 8 retinas, *p < 0.001; t-test).

Anatomical studies show that inputs to proximal starburst dendrites originate mainly from BC7s, while inputs to distal dendrites arise from BC5s (including types BC5i, o & t; Greene et al., 2016; Ding et al., 2016), indicating that the kinetic differences of iGluSnFR responses may reflect the properties of distinct BC types. By using AAV-8BP/2 vector containing CAG promotor, we directly monitored glutamate release at BC7s axon terminals (Matsumoto et al., 2021; **Fig. 3**). We identified axon terminals of BC7s based on the depth of ON starburst cell dendrites that were genetically labeled by tdTomato (Matsumoto et al., 2021). We found that iGluSnFR responses at BC7 terminals were sustained. The most appreciable changes in iGluSnFR fluorescence occurred at the axon terminals, suggesting that the sensor signals reflect the vesicle release dynamics of individual BC7s (James et al., 2019). The peak/plateau ratio was independent of response amplitude, indicating the degree of sustained is not exaggerated by signal/noise issues (**Fig. S1**). Unfortunately, we failed to devise ways to express the sensor in other BC5s and could not directly confirm that these were the sources of transient signals observed in distal starburst dendrites. However, the results lend support to the idea that BC7s are the source of sustained proximal input.

**Fig. 3:**
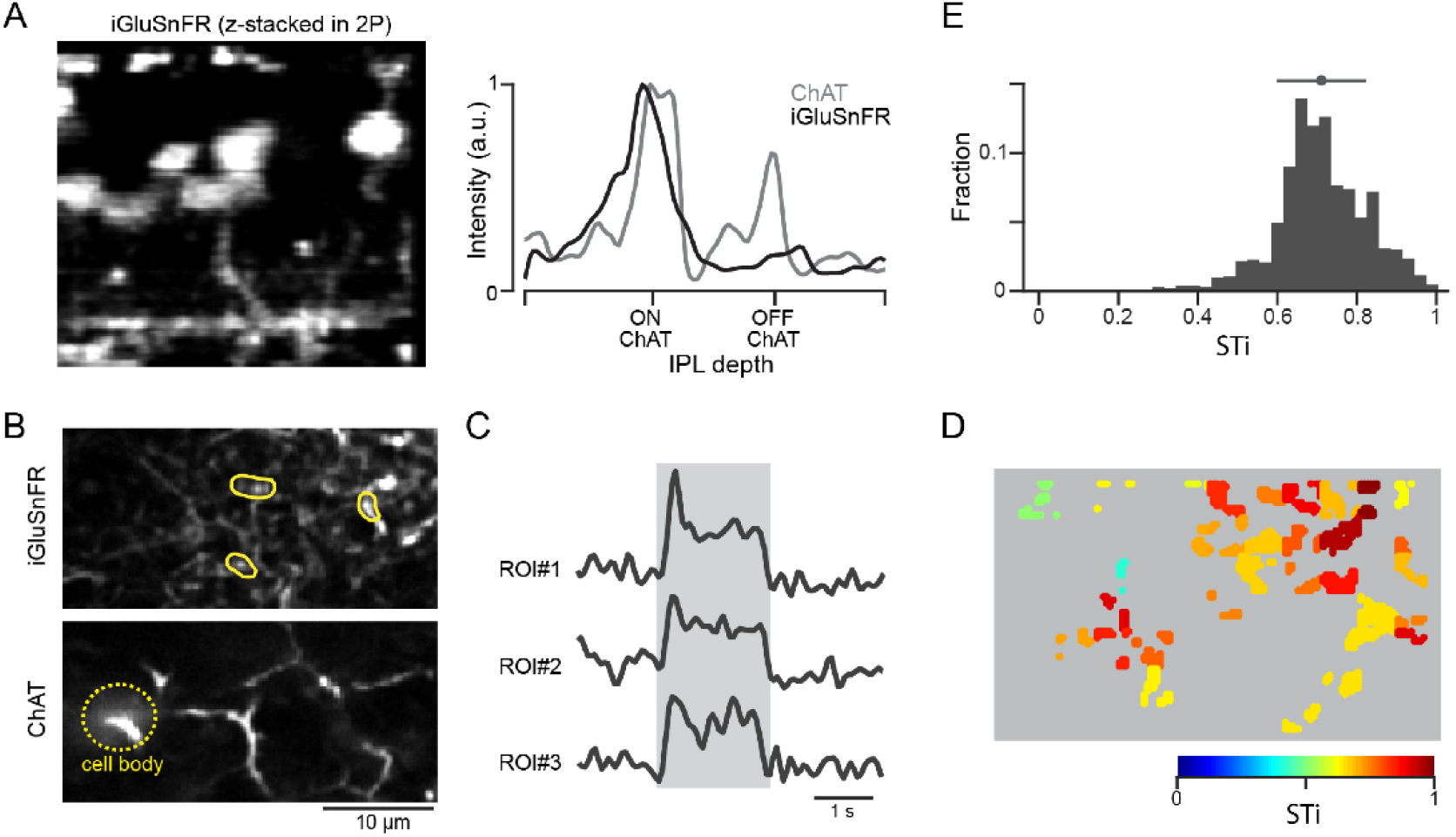
Expressing iGluSnFR in BC7 axon terminals reveals their sustained output. (A) Cross-section of an image stack showing iGluSnFR labelled BC7 (left). The intensity profiles of the BC terminals labelled with iGluSnFR (grey) and starburst dendrites labelled with tdTomato (black) across the inner plexiform layer (IPL) show that BC terminals co-stratify with dendrites of ON starbursts (right). (B) iGluSnFR expression in BC7 terminals (top) imaged at the same depth as the proximal ON starburst dendrites labelled with tdTomato (bottom) (C) Light-evoked glutamate signals (right) extracted from three ROIs shown in B (left). The grey band indicates the stimulus duration. (D) Heat maps of the STis for all identified ROIs. (E) A histogram of STis for the light-evoked responses for all ROIs. Top, mean (circle), and s.d. (horizontal bar) of the STis.

In contrast to spots of light, when white-noise stimuli were to characterize the temporal properties of BCs using reverse-correlation techniques, we failed to observe significant kinetics differences in proximal and distal iGluSnFR responses. Similar to results form a recent electrophysiological study (Fransen and Borghuis, 2017), we found the input impulse responses were biphasic (**Fig. S2**). Thus, the probability of glutamate release from BC terminals appears to be transiently depressed, following a burst of vesicle release, during continuous stimulus regimes. However, biphasic kinetics were observed for proximal and distal dendritic responses, consistent with parallel imaging studies (Franke et al., 2017; Strauss et al., 2021). As the biphasic nature of the distal—but not proximal—bipolar cell kinetics is critical to the success of models generating direction selectivity (Kim et al., 2014; Fransen and Borghuis, 2017), we conclude that under conditions where the circuit is continually stimulated, input kinetics are unlikely to play a role in shaping direction selectivity for objects moving smoothly across the starbursts’ receptive field.

### BC output kinetic differences are shaped largely by excitatory network mechanisms

Next, to investigate whether cell-intrinsic or network mechanisms shape BC kinetics, we examined how iGluSnFR responses were affected by stimulus size. Increasing the spot diameter systematically decreased the peak amplitude of BC responses, indicative of the recruitment of the inhibitory surround (**Fig. 4A**; Franke et al., 2017). Importantly, the distinction in the kinetics of proximal and distal inputs remained clear across stimulus sizes, although they were generally more pronounced for stimuli that were > 200 μm (**Fig. 4B**, control; n=71 proximal and 431 distal ROIs, 5 FOVs, 4 retinas; *p<0.001, Kolmogorov-Smirnov test). Indeed, the application of a cocktail of antagonists containing both GABA and glutamate receptor antagonists (5 μM gabazine and 100 μM TPMPA, 20 μM CNQX, respectively)—which blocks inhibitory inputs from amacrine and horizontal cells—augmented responses, especially those evoked by larger stimuli (**Fig. 4A**). These effects reduced the overall STi as compared to control, but the kinetics of the iGluSnFR responses at proximal and distal ROIs remained distinct **(Fig. 4B**, drug cocktail; n=71 proximal and 431 distal ROIs, 5 FOVs, 4 retinas; *p<0.001, Kolmogorov-Smirnov test). Thus, while the inhibitory networks appear to modulate BC responses, they do not appear to be required for the generation of sustained/transient differences observed here.

**Fig. 4:**
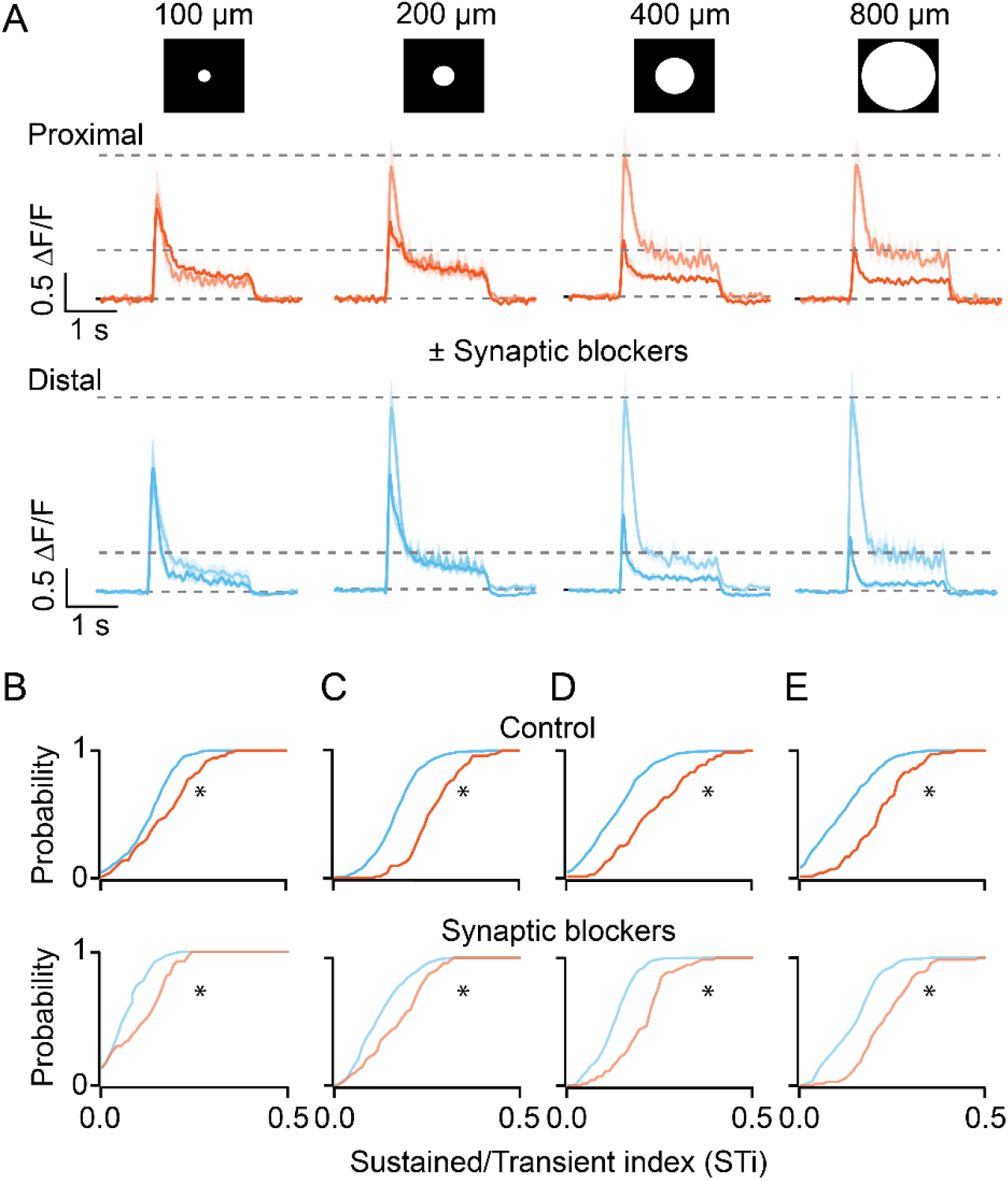
Kinetic differences in iGluSnFR signals are apparent across a range of stimulus sizes and persist in the presence of inhibitory receptor blockers. (A) The average iGluSnFR signals were evoked by spots of different diameters (100 μm - 800 μm). Responses were averaged across five proximal (orange) or distal (blue) FOVs. Responses measured under control (dark traces) conditions and in the presence of synaptic blockers (SR, TPMPA, and CNQX; light traces) are overlaid. Shading indicates ± s.e.m. (B-E) Cumulative distributions of STis for ROIs in the proximal and distal FOVs under control and blocker conditions for different stimulus sizes. n = 5 FOVs, 4 retinas, *p < 0.001; Kolmogorov-Smirnov test.

In addition to demonstrating the inhibitory effects of the surround, blocking glutamate/GABA/glycine receptor mediated pathways using the drug cocktail also revealed a lateral excitatory pathway. Lateral excitation manifests in the plateau but not the peak phase of the response (**Fig. 4A**). In contrast to the peak phase, which could optimally be triggered by small spots (∼100 μm diameter), the plateau phase continued to grow with spot size. It reached a maximum when the spot was ∼200 μm in diameter, which is significantly larger than the dendritic field of BCs (**Fig. 4A**). Such lateral excitation has been attributed to electrical coupling, which occurs between BCs as well as amacrine cells (Sigulinsky et al., 2020; Arai et al., 2010; Asari and Meister, 2014). The finding that the kinetic diversity of BC responses is maintained in the absence of inhibition suggests it is largely shaped by excitatory networks.

### BC output kinetics contribute to direction selectivity

Next, we tested how the apparent kinetic diversity in BC input impacts direction selectivity. Since the BC kinetics inferred from iGluSnFR measurements are in part dictated by the properties of the indicator, it remains unclear how they relate to the starbursts’ physiological responses mediated by endogenous AMPA receptors. To address this issue, we first used optical deconvolution methods to estimate the time-varying vesicle release rates from individual BCs, and then used a computational model to understand how trains of vesicles are transformed into AMPA receptor-driven voltage signals in starburst dendrites.

Time-varying vesicle release rates from proximal and distal BCs were estimated by deconvolving the iGluSnFR response with an idealized quantal response described by an alpha function with decay kinetics ∼30 ms (**Fig. 5B**, inset), which was obtained by matching the kinetics of ‘spontaneous’ quantal responses (**Fig. 5A**). The resulting release rates were discretized using a Poisson process, yielding an estimate of the temporal pattern of single vesicle release events (**Fig. 5C**). For this analysis, the quantal size for each ROI was estimated using fluctuation analysis (see Methods). Indeed, convolving the average quantal response with the inferred trains of vesicle release events (**Fig. 5C**) produced a signal similar to the original iGluSnFR measurement (**Fig. 5D**). The release profiles of proximal and distal BCs obtained in this manner support the notion that they have different capacities to release vesicles in response to light stimuli: proximal BCs release vesicles throughout the duration of the stimulus, while distal BCs release a large fraction of their vesicles at the onset of the light stimulus (**Fig. 5E**).

**Fig. 5:**
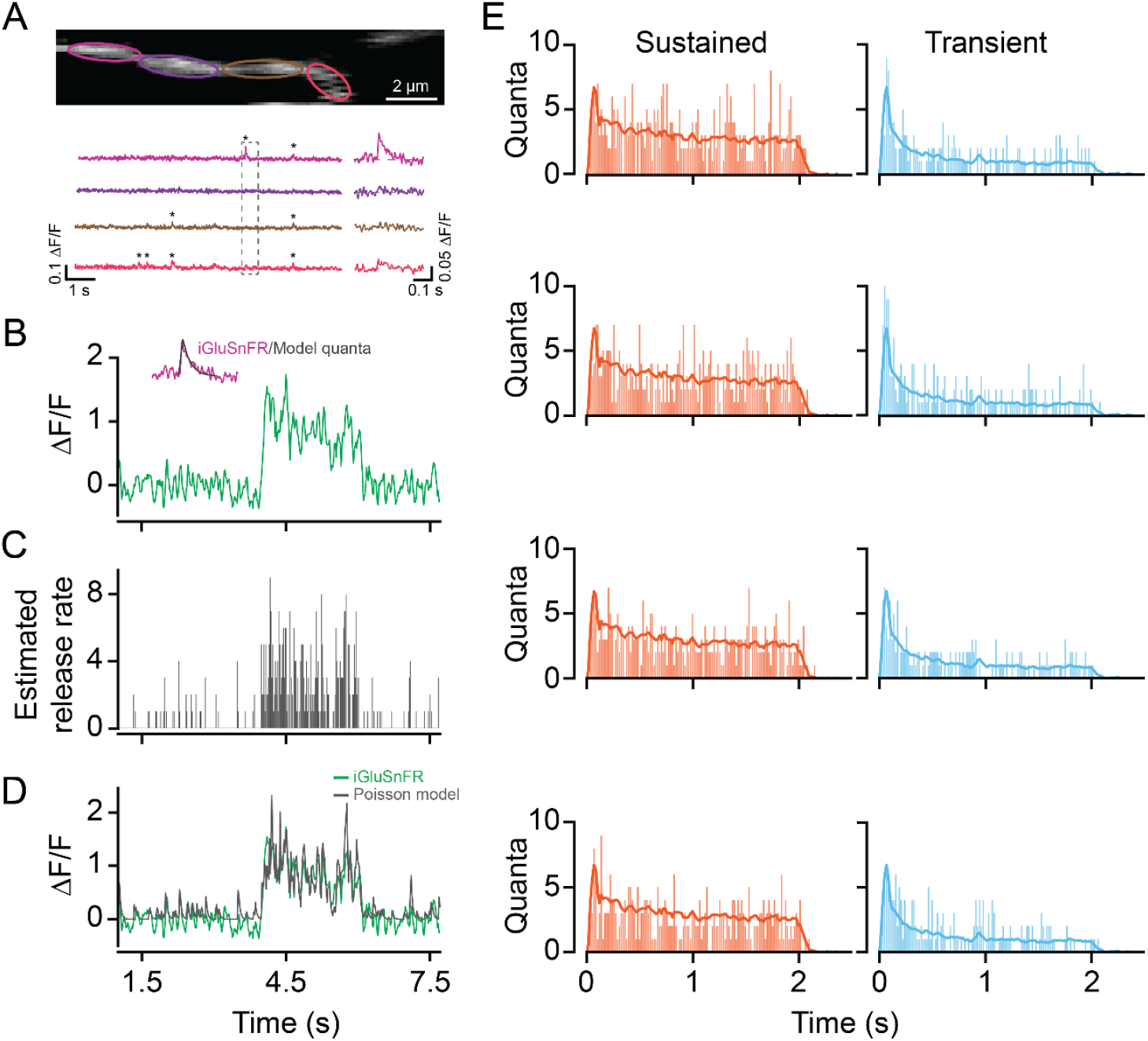
Time-varying vesicle release rates estimated using temporal deconvolution. (A) Spontaneous iGluSnFR signals measured in neighboring ROIs across a small dendritic section (color-coded to match ROIs). (B) A typical light-evoked iGluSnFR signal measured from proximal dendrites; inset: iGluSnFR quantal event fitted with an alpha function. (C) A time-varying release rate was estimated by deconvolving the iGluSnFR signal with the quantal signal (shown in B). (D) Convolving the estimated release rate with a unitary event recapitulates the shape of the original iGluSnFR response. (E) Example vesicle release rates for sustained (orange) and transient (blue) iGluSnFR responses. Solid line indicates average vesicle release rates for all ROIs. n = 50 each, proximal and distal ROIs.

To understand how vesicle release dynamics from BCs shape starburst responses we constructed a simple multi-compartmental model (NEURON) using previously described cable properties of starbursts (Tukker et al., 2004; Vlasits et al., 2016). Sustained and transient BCs were assumed to reflect the properties of BC7 and BC5s, respectively, and distributed non-uniformly across the starburst dendrite in accordance with the anatomical data (Ding et al., 2016; Greene et al., 2016, **Fig. 6A**; also see Methods). Simulated moving edges ‘activated’ BCs in succession, triggering streams of AMPA receptor-mediated miniature-like events (|_decay_ ∼ 0.54 ms; Vlasits et al., 2016) at different points along the starburst dendrite. The voltage responses measured from the model cell soma were qualitatively like those measured experimentally. For example, responses in the preferred direction rose rapidly compared to those evoked in the null direction, owing to the asymmetric distribution of inputs (**Fig. 6A** middle panel; Ankri et al., 2020).

**Fig. 6:**
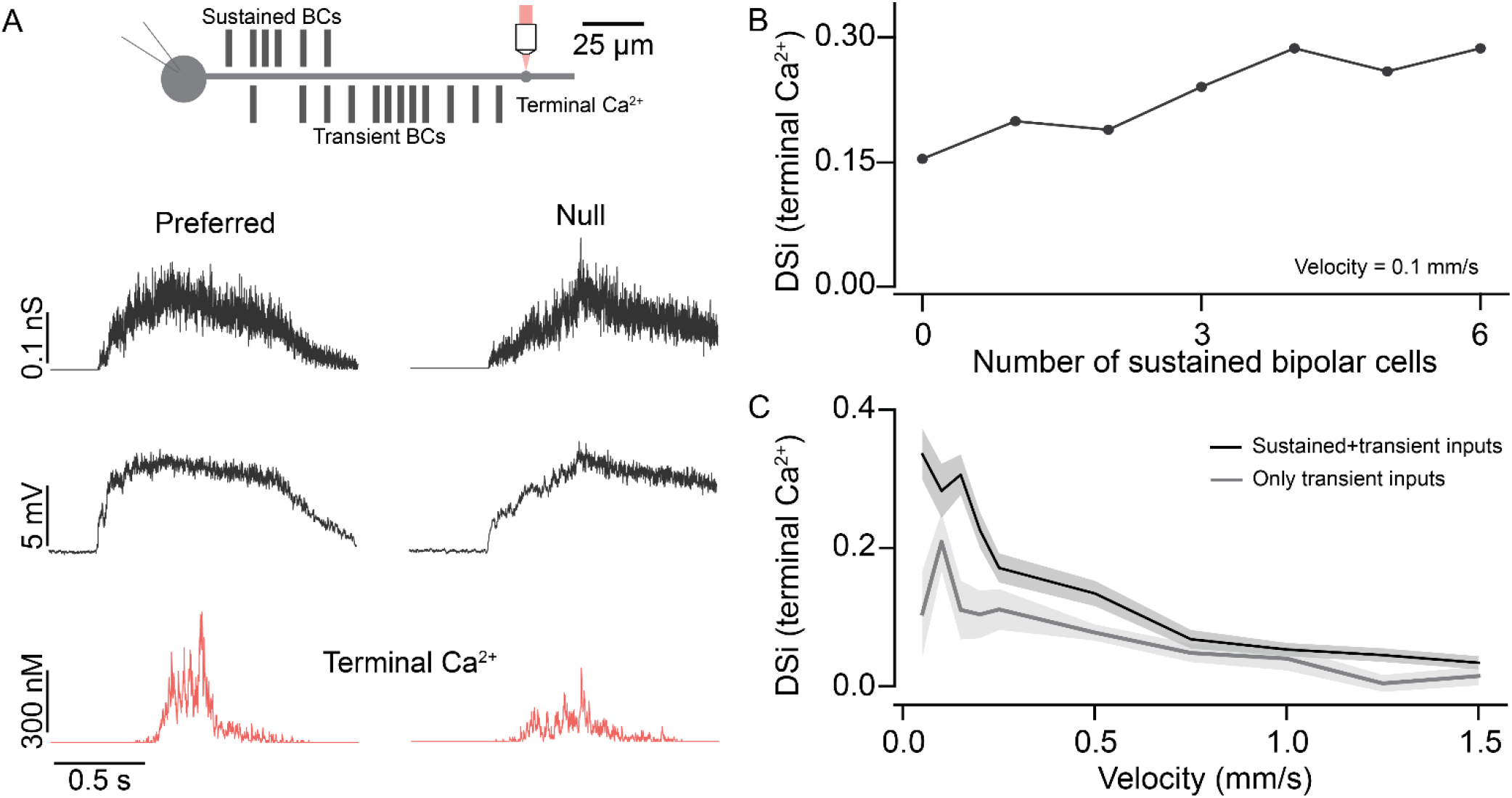
Input kinetics shape direction selectivity at low stimulus velocities. (A) Schematic representation of locations of somatic voltage and terminal Ca^2+^ recordings from a model SAC under simulated conditions (top). Bipolar cell conductances, somatic voltage, and terminal Ca^2+^ responses (bottom) measured in the preferred and null direction from the model SAC when simulated using moving edges. (B) Direction selectivity index (DSI) of peak Ca^2+^ (terminal) responses vs number of inputs from sustained BCs. (C) Direction selectivity index (DSI) of peak Ca^2+^ (terminal) responses vs velocity.

Notably, the distinct temporal kinetics of vesicle release from the distinct types of BCs appeared to play a critical role in the generation of direction selectivity. When both sustained and transient inputs were used to drive starburst dendrites, robust direction selectivity was observed. This was most pronounced in the dendritic intracellular Ca^2+^ signals (**Fig. 6A**, bottom traces), likely due to the non-linear properties of Ca^2+^ channels (Tukker et al., 2004). These model data accurately recapitulate direction selectivity measured using two-photon Ca^2+^ imaging (Euler et al., 2002). However, when the proximal (sustained) BCs were incrementally converted into transient ones, starting from the most distal and proceeding inward until all inputs were made transient, direction selectivity decreased linearly (**Fig. 6B**). This is presumably because the amount of asymmetric temporal summation decreases as inputs with weaker plateau phases are swapped in. Finally, as expected for a mechanism that relies on a fixed delay, direction selectivity was strongly dependent on stimulus velocity, being most robust for stimuli moving less than 300 μm/s (**Fig. 6C**). Thus, we conclude that in the context of realistic estimates of glutamate input kinetics, the ‘space-time’ wiring model for direction selectivity holds merit under specific stimulus conditions.

## Discussion

Direction selectivity in starburst dendrites is one of the most striking computations observed in the retina. Despite being intensely investigated over the last two decades, a core mechanisms that underlie direction selectivity in starbursts have remained elusive. This is because a variety of overlapping synaptic mechanisms may contribute to direction selectivity, enabling the computation to occur robustly across a range of stimulus parameters including stimulus speed, contrast, and size. In this study, we determined the kinetics of glutamate input across individual starburst dendrites and propose that they play an important role in shaping direction selectivity for slow-moving objects, providing support for the ‘space-time’ wiring mechanism for DS.

A recent flurry of glutamate imaging studies has provided us with unique insights into the functional diversity of BCs (Borghuis et al., 2013; Franke et al., 2017; Gaynes et al., 2021; Matsumoto et al., 2021; Park et al., 2014; Strauss et al., 2021; Yonehara et al., 2013). Large-scale surveys, in which glutamate release was measured from a large number of BC axon terminals across the entire IPL, demonstrated that local responses across all BC types of the same response polarity are similar in their kinetics. Our current study supports and extends the result from the previous studies in several important ways. First, previous studies distinguished between the different BC types using elegant algorithms that clustered cells based on their response properties as well as their axonal stratification depth in the IPL. While this method provides unique insights into the general organization of glutamate signaling in the inner retina, it falls short in unambiguously delineating BC types in the starburst circuit that have overlapping axonal stratification patterns (Franke et al., 2017; Strauss et al., 2021; Gaynes et al., 2021). Here, by restricting iGluSnFR expression to the starbursts, and imaging along individual dendrites and/or at different depths within the starburst dendritic layers, we obtained estimates of the kinetics of proximal and distal input to starbursts, without explicitly attempting to pinpoint their specific sources.

Clear kinetic differences between proximal and distal regions were observed for spots of varying sizes (>100 μm in diameter). However, when small spots that just covered the BC receptive field were utilized, the proximal and distal responses had similar kinetics, similar to previous reports (Franke et al., 2017). Moreover, kinetic differences were discernable for stationary or slow-moving spots. When a Gaussian white noise (10 or 20 Hz) stimulus was used, events measured from proximal and distal dendritic regions were similar. Together, these results suggest the possibility that BC kinetics play a role in starburst dendritic computations under specific stimulus conditions.

The finding that kinetic differences manifest for spot sizes larger than BC dendritic fields implies that network mechanisms are involved in shaping BC kinetics (**Fig. 4A**). Previous studies have suggested that the functional diversification of BC responses arises mainly from GABAergic and/or glycinergic inhibition mediated by amacrine cells (Franke et al., 2017; Franke and Baden, 2017). Thus, it was surprising to observe clear kinetic differences when all inhibitory pathways were blocked. While the kinetic differences could be influenced by inhibition, inhibition was not required to generate kinetic differences.

At first pass, this indicates that kinetic differences arise from cell-intrinsic factors (Awatramani and Slaughter, 2000). However, since the size of the spot that evoked an optimal response was significantly larger than BC dendritic fields suggests that the lateral excitatory pathway contributes to BC responses. Specific types of BCs are known to be coupled to each other as well as to select amacrine cells including the AII and A8 amacrine cells (Arai et al., 2010; Asari and Meister, 2014; Demb and Singer, 2012; Sigulinsky et al., 2020; Yadav et al., 2019; Zhang and Wu, 2009). Such coupling has been implicated in extending BC receptive fields spatially in salamander and fish retina (Hare and Owen, 1990; Saito and Kujiraoka, 1988), and in mice, the electrical coupling can non-linearly affect the output of individual BCs (Kuo et al., 2016). Additionally, the finding that kinetic differences persist when rod-mediated signals are blocked by CNQX suggests that kinetic differences cannot be attributed solely to changes associated with rod/cone vision (Awatramani and Slaughter, 2000). Thus, BC receptive field structures may be influenced in more complex ways than earlier envisioned, being influenced both by inhibitory and excitatory synapses (Franke et al., 2017).

Another major advance made here is to relate iGluSnFR signals to endogenous vesicle release rates from BC terminals and eventually to starburst function. As the expression pattern, affinity, and kinetic properties of iGluSnFR are vastly different from glutamate receptors that mediate synaptic responses, interpreting iGluSnFR signals is not straightforward. Previous studies have shown iGluSnFR signals reflect local glutamate release from individual BC terminals, rather than global “spill-over” signals, despite being expressed uniformly along starburst dendrites (Borghuis et al., 2013; Franke et al., 2017; Gaynes et al., 2021; Matsumoto et al., 2021; Strauss et al., 2021). Here, we were also able to measure spontaneous ‘quantal-like’ events using high-speed imaging. This enabled us to use temporal deconvolution methods to assess release rates at proximal and distal sites. Assuming the light-evoked responses were made up of a linear supposition of quantal events, we estimate the peak instantaneous release rate to be 5-10 vesicles/s, which is in line with previous *in vivo* estimates made in zebrafish BC terminals (James et al., 2021, 2019). We also estimated the steady-state release rates to be ∼ 3 vesicles/s at proximal sites and ∼1 vesicle/s at distal sites, which is well below the maximum release rate estimated for photoreceptor synapses, which also contain ribbons (Hays et al., 2021, 2020).

To understand how BC kinetics impact starburst function, previous studies have mainly relied on phenomenological models such as the linear-non-linear cascade, in which responses are estimated by successively passing the input stimulus through a linear temporal filter followed by a static non-linear transformation (Kim et al., 2014; Franke et al., 2017; Strauss et al., 2021; Fransen and Borghuis, 2017). These simplified models leave out the biophysical details of the starburst, and the extent to which they reproduce the input-output function of real starburst dendrites remains unclear. This issue was addressed to some extent by detailed compartmental modeling of the starburst circuit (Ezra-Tsur et al., 2021). However, in this study, the key model parameters, including the vesicle release rates and the specific distribution of BC types along starburst dendrites, were determined by a complex algorithm that optimizes direction selectivity at relatively fast speeds (900 μm/s). By contrast, we demonstrate that in a simplified circuit model that is constrained by experimentally determined vesicle release rates and by the spatial distributions of sustained and transient input to starburst dendrites, differences in input kinetics mainly enhances direction selectivity at low velocities (< 300-400 μm/s). Thus, the robust direction selectivity that is experimentally observed in starbursts, across the range of velocities (100-2000 μm/s; Ding et al., 2016), must rely on additional mechanisms. Further work is needed to establish how the excitatory and inhibitory network mechanisms, previously described, work together to produce direction selectivity that is invariant across a large range of velocities.

## Methods

Animals: Experiments were performed using ChAT-IRES-Cre (Δneo) mice on a C57BL/6J background (Jackson laboratory, #031661). Mice were P21 or older and of either sex. Animals were housed under 12 hour light/dark cycles. All procedures were performed in accordance with the Canadian Council on Animal Care and approved by the University of Victoria’s Animal Care Committee, or in accordance to Danish standard ethical guidelines and were approved by the Danish National Animal Experiment Committee (Permission No. 2015-15-0201-00541; 2020-15-0201-00452).

Viral injections: For intravitreal injections, mice were anesthetized by administering isoflurane (2-3 % at 1-1.5 L/min; Fresenius Kabi Canada Ltd.) mixed with oxygen (1-3 %) through a vaporizer. Buprenorphine was administered subcutaneously (0.05-0.1 mg/kg body weight) as an analgesic. After creating a small hole at the margin of the sclera and cornea with a 30-gauge needle, a volume of 1-1.2 μl of the viral plasmid pAAV.hSyn.Flex.iGluSnFR.WPRE.SV40 (gift from Dr. Loren Looger; Addgene plasmid # 98931; http://n2t.net/addgene:98931; RRID:Addgene_98931) was injected into the vitreous humor of either the left or right eye using a Hamilton syringe (syringe: 7633-01, needle: 7803-05, point style 3, length 10 mm). Mice were then returned to their home cage, placed on a heating pad, and monitored until fully recovered. Imaging experiments were performed at least three weeks after injections.

Tissue preparation: Mice were dark-adapted for at least 60 minutes before being anesthetized with isoflurane (Fresenius Kabi Canada Ltd.) and decapitated. Retinas were extracted in Ringer’s solution under a dissecting microscope equipped with infrared optics. The isolated retina was then whole-mounted onto a glass coverslip coated with poly-L-lysine (Sigma Aldrich). The mounted retina was then placed into a recording chamber under the microscope and perfused continuously with oxygenated Ringer’s solution (95% O_2_/5% CO_2_; 35°C). Retinas were visualized with a Spot RT3 CCD camera (Diagnostics Instruments) through a 40x water-immersion objective on a BX-51 WI microscope (Olympus Canada).

Two-photon imaging: For iGluSnFR imaging, we used an Insight DeepSee^+^ laser (Spectra Physics) tuned to 920 nm, which was guided by an 8 KHz resonant-galvo-galvo scan head (Vidrio) for two-photon excitation. The projector LEDs were coordinated to only be on during the turn-around phase of the resonant mirror, thereby preventing the bright projector light from contaminating fluorescence signals. Green and red fluorescence emissions were detected using two photomultiplier tubes (PMTs; Hamamatsu) that were equipped with appropriate bandpass filters (525/45 and 625/90; Semrock). A high-speed current amplifier (200 MHz, Edmund Optics) was used to count individual photon events from the PMTs, further reducing acquisition noise. Images were acquired using ScanImage software (Pologruto et al., 2003; Vidrio Technology). While most recordings from proximal and distal dendritic regions were acquired consecutively at a frame rate of 58.25 Hz (256 × 256 pixels), in some experiments the responses from the two regions were recorded near-simultaneously at a frame rate of 22.5 Hz (256 × 256 pixels) using an electrically-tunable lens (ETL; EL-10-30-TC-NIR-12D, Optotune; see methods Murphy-Baum and Awatramani, 2022). Data using either method were compiled in Figure 2C-D.

Visual stimulation: Visual stimuli were produced using a digital light projector, and were focused onto the photoreceptor layer of the retina through the sub-stage condenser. The background illuminance was measured to be ∼ 1000 photon/μm^2^/s. Visual stimuli were generated and presented using StimGen, a python-based visual stimulation interface (https://github.com/benmurphybaum/StimGen). For all experiments, static flashes of different spot sizes (100, 200, 400, and 800 μm) were presented for a duration of 2 s, with an initial delay of 4 s.

Pharmacology: The following concentrations (in μM) of antagonists were used for experiments: 20 CNQX disodium salt (Hello Bio, U.K.); 100 TPMPA (Tocris Bioscience), and 5 SR-95531 (Hello Bio, U.K.). Antagonists were initially prepared as stock solutions in distilled water, or in DMSO in the case of CNQX. During experiments, drugs were freshly prepared froma stock solution in carboxygenated Ringer’s solution. All drug solutions were bath applied to the tissue for at least 5 minutes prior to recording.

Quantification and statistical analysis: All imaging analysis and statistical comparisons were performed in Igor Pro (WaveMetrics). Fluorescence signals from either 5×5 μm or 10×10 μm regions of interest (ROI) were calculated as ΔF/F, and smoothed using a 2^nd^ order Savitsky-Golay filter before further analysis. Signal-to-noise ratio (SNR) was computed for each ROI as:

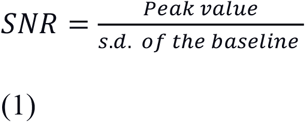

where baseline fluorescence was measured as the fluorescence signal in a 1 s window before the start of the stimulus. Only ROIs with responses above 4 to 5 SNR were selected for further analysis.

The sustained transient index (STi) for each ROI was calculated as:

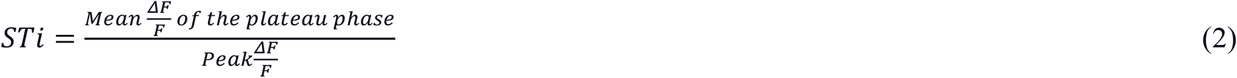

where the mean plateau ΔF/F is calculated for the last 1 s of the stimulus duration. Statistical tests and sample sizes are indicated in the figure legends.

Computational modeling: Representative sustained and transient bipolar cell release rates were calculated via an inverse fourier transform method (Van der Kloot, 1988; Diamond & Jahr, 1995; Awatramani et al. 2007), using an iGluSnFR quantal template (2 ms rise, 30 ms decay) scaled by the quantal size estimate (QSE) of steady-state (plateau phase) iGluSnFR responses to static spot presentation (Katz and Miledi, 1972; Awatramani et al 2007). Averages responses from proximal and distal scan fields were used to calculate the prototypical sustained and transient bipolar cell release rates, respectively (n = 50 ROIs, each proximal and distal site, 6 retinas, 7 FOVs).

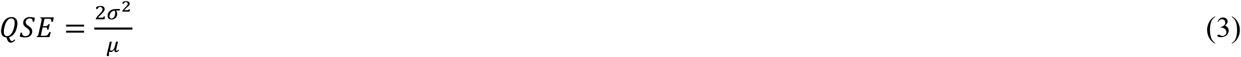

where *σ* and *μ* are the variance and the mean of the steady-state of iGluSnFR responses.

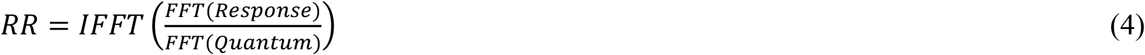

where *RR* is the release rate, *Response* is the iGluSnFR response evoked by a spot of light and *Quantum* refers to the spontaneous quantal iGluSnFR signal.

A simple ball and stick compartmental model was constructed in the NEURON simulation environment, with one somatic compartment and three dendritic compartments (initial, middle, and terminal dendrites). Input to this isolated SAC computational unit was driven by model bipolar cell synapses distributed according to previous electron microscopy measurements (Ding et al. 2016). Specifically, the proximally-located, sustained bipolar cells were positioned according to the measured distribution of BC7 contacts. Distally-located, transient bipolar cells were positioned according to the measured distribution of BC5 contacts, which includes all subtypes of BC5. These bipolar cell inputs were then triggered by a simulated light edge moving over a range of velocities (0.1 to 2 mm/s). Thus, each bipolar cell input began releasing glutamate according to its location relative to the stimulus. For each bipolar cell synapse, on each stimulus presentation, the prototypical release rates were translated into discretized quantal trains via a Poisson process (sampled at model time step of 1 ms), and applied to the SAC dendrite as a series of AMPAR conductance events (rise 0.14 ms, decay 0.54 ms, reversal 0 mV, max conductance 156.5 pS (scaled down with increasing velocity according to iGluSnfr recordings). The somatic membrane voltage and terminal dendritic Ca^2+^ accumulation was recorded to assess post-synaptic activation. The direction-selective index (DSI) of the model was assessed by subtraction of the peak terminal Ca^2+^ concentration in centripetal stimulation from that of centrifugal stimulation.

### Model Parameters

Model Parameters:
|  | Soma | Initial dendrite | Middle dendrite | Terminal dendrite |
| --- | --- | --- | --- | --- |
| Length [$\mu\text{m}$] | 7 | 10 | 120 | 20 |
| Diameter [$\mu\text{m}$] | 7 | 0.4 | 0.2 | 0.2 |
| Ra [$\Lambda\oplus\text{cm}$] | 150 | 150 | 150 | 150 |
| gleak (hh) [$\text{S}/\text{cm}^2$] | 4.6e-5 | 4.6e-5 | 4.6e-5 | 4.6e-5 |
| eleak (hh) [$\text{S}/\text{cm}^2$] | 0 | 0 | 0 | 0 |
| $\text{Na}_v$ (hh) [$\text{S}/\text{cm}^2$] | 0 | 0 | 0 | 0.0001 |
| $\text{K}_v$ [(hh) [$\text{S}/\text{cm}^2$] | 0.01 | 0 | 0 | 0 |
| $\text{K}_m$ (hh) [$\text{S}/\text{cm}^2$] | 0 | 0 | 0 | 0 |
| N-type $\text{Ca}^{2+}$ [$\text{S}/\text{cm}^2$] | 0 | 0 | 0 | 0.0003 |
| P-type $\text{Ca}^{2+}$ [$\text{S}/\text{cm}^2$] | 0 | 0 | 0 | 0.0001 |
| $\text{K}_{v3.3}$ | 0 | 0 | 0 | 0.05 |

### Bipolar Cell Locations

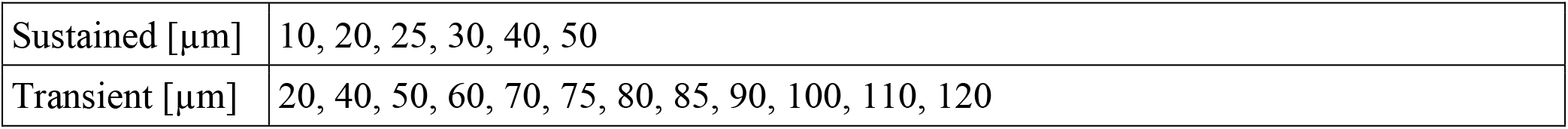

Bipolar Cell Locations:
|  |  |
| --- | --- |
| Sustained [$\mu\text{m}$] | 10, 20, 25, 30, 40, 50 |
| Transient [$\mu\text{m}$] | 20, 40, 50, 60, 70, 75, 80, 85, 90, 100, 110, 120 |

## Figures and Legends

**Supplementary Fig. 1:**
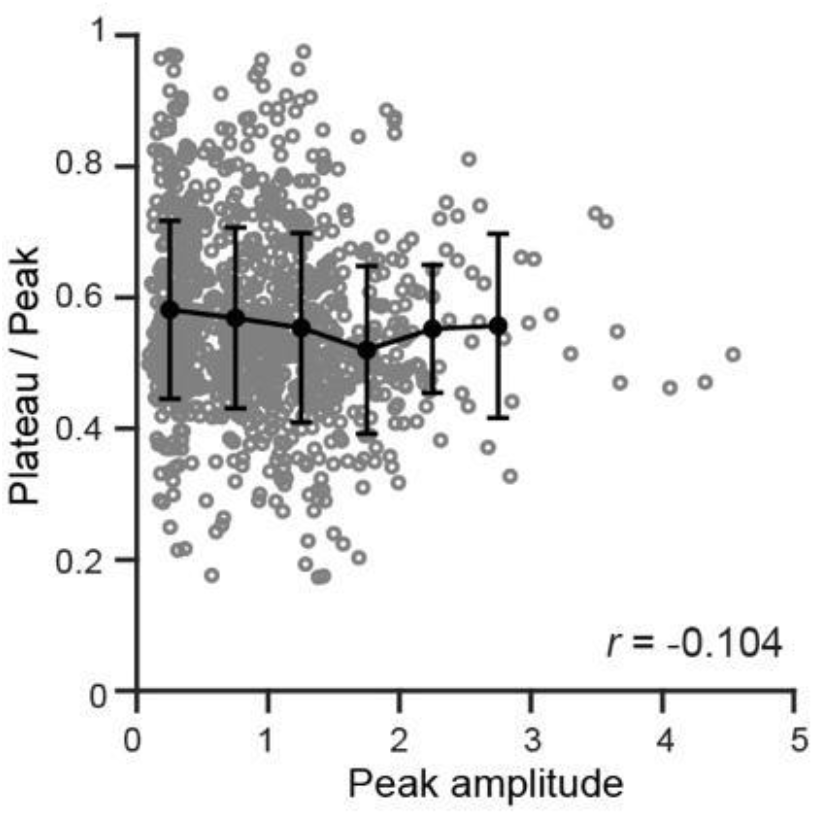
Temporal properties of BC7s remain stable across response amplitude.

**Supplementary Fig. 2:**
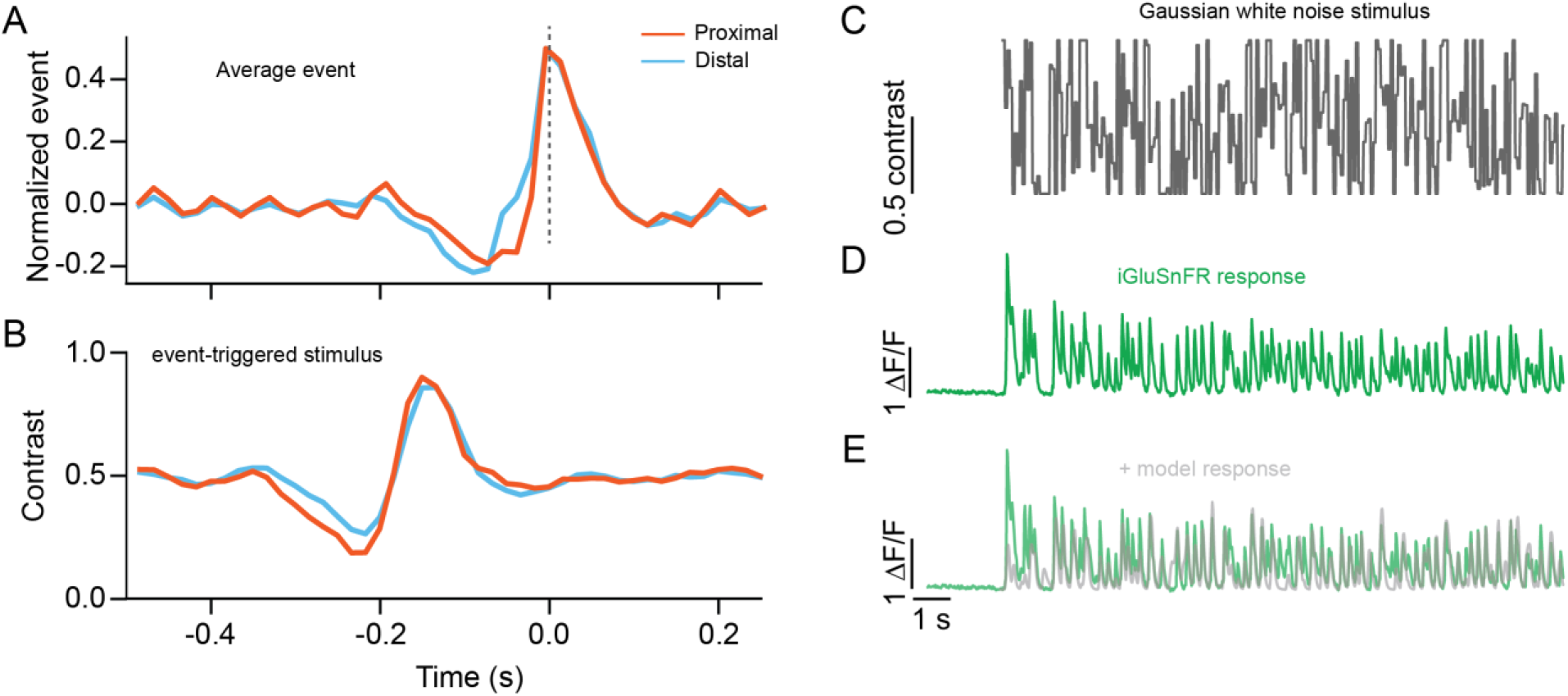
Temporal properties of proximal and distal input revealed by noise analysis. (A) Average waveform of iGluSnFR responses to Gaussian white noise measured in the proximal or distal dendritic regions of starbursts. (B) Event-triggered average stimulus waveform, which represents the BCs’ preferred stimulus, is shown for proximal and distal regions. (C) The intensity profile of the 20 Hz Gaussian white noise stimulus. (D) The iGluSnFR response to the stimulus in C. (E) Model iGluSnFR response obtained by convolving stimulus with linear filter shown in B

